# Systemic HIV-1 Infection Produces a Unique Glial Footprint in Humanized Mouse Brains

**DOI:** 10.1101/178285

**Authors:** Weizhe Li, Santhi Gorantla, Howard E. Gendelman, Larisa Y. Poluektova

## Abstract

Studies of innate glial cell responses for progressive human immunodeficiency virus type one (HIV-1) infection are hindered by the availability of relevant small-animal models. To overcome this hindrance, a mouse model reconstituted with humanized brain and immune systems was created. Newborn NOD/SCID/IL2Rγc^-/-^ mice of both sexes were transplanted with human neuroglial progenitors (NPC) and hematopoietic stem cells. Intraventricular injection of NPC yielded an anatomical symmetrical glia (human astrocyte and oligodendrocyte) repopulation of the mouse brain. The human glia were observed in periventricular areas, white matter tracts, the olfactory bulb and brain stem. HIV-1 infection of these dual humanized mice led to meningeal and perivascular human leukocyte infiltration into brain. The species-specific viral-neuroimmune interactions in the infected animals were identified by deep RNA sequencing. In the corpus callosum and hippocampus overlapping human-specific transcriptional alterations were seen for interferon type 1 and 2 signaling pathways (*STAT1, 2, IRF9, ISG15, IFI6*) and a range of host antiviral responses (*MX1, OAS1, RSAD2, BST2, SAMHD1)* in infected animals. Glial cytoskeleton reorganization, oligodendrocyte differentiation and myelin ensheathment (*MBP, MOBP, PLP1, MAG* and *ZNF488*) were downregulated. The data sets were confirmed by real-time PCR. The viral defense signaling patterns observed in these mice parallels the neuroimmune communication networks present in the HIV-1 infected human brain. In this manner, the new mouse model can facilitate discovery of therapeutics, viral eradication targets for virus induced nervous system diseases, and simplify HIVCure research approaches.

**Summary Statement:** We created mice with both a humanized brain and an immune system. The animals were used to investigate glial responses to HIV-1 infection. At a transcriptional level we defined the interactions between human glia and immune cells in the presence of the systemic HIV-1 infection. Noticeably, altered transcriptional changes were human specific. At five weeks after viral infection humanized mouse brain displayed potent interferon-mediated antiviral innate immune responses and alteration of neuronal progenitors differentiation and myelination. This model can be used to tests both diagnostic and therapeutic interventions for cure HIV-associated brain impairment.

## Introduction

Despite sustained investigation of the pathophysiological mechanisms of human immunodeficiency virus (HIV)-associated neurocognitive disorders (HAND), virus-induced changes in neuroimmune communications are incompletely understood (Gelman et al., 2012; Saylor et al., 2016). As the virus cannot infect rodent cells co-morbid disease conditions that lead to impaired immune, glial, and neural cell function have not yet been reflected in a small animal. Transgenic mice containing the whole viral genome or individually expressed proteins and animals injected with viral proteins or human infected cells in brain tissue can only mirror disease components (Gorantla et al., 2012). Lentiviral infections of non-human primates, cats, and ungulates can mirror HIV nervous system comorbidities (Narayan and Clements, 1989) but only some of the natural infectious processes (Jaeger and Nath, 2012). Such limitations are also operative in mice possessing a sustainable human immune system. While humanized animals are susceptible to HIV infection and support persistent viral replication the resultant structural and functional changes that occur in the brain are limited and not always reflective of human disease (Boska et al., 2014; Dash et al., 2011; Gorantla et al., 2010). The absence of human glial cells and differences in viral responses in the setting of divergent patterns of murine and human brains limit mechanistic studies (Oberheim et al., 2009).

Astrocytes and oligodendrocytes, the most abundant cell type in the human central nervous system, play pivotal roles in brain homeostasis and metabolism. Astrocytes not oligodendrocytes show limited susceptibility to HIV infection (Chauhan, 2015; Farina et al., 2007; Pfefferkorn et al., 2016; Russell et al., 2017). During disease, communication between astrocytes and oligodendrocytes and infected mononuclear phagocytes (MP; monocyte macrophages and microglia) herald changes in the brain’s pro-inflammatory microenvironment (Bednar et al., 2015). The restriction to viral infection reflect absent receptors for viral entry (Schweighardt and Atwood, 2001) and coordinate with less studied intracellular innate immunity protective pathways. Each do not permit the viral life cycle to reach completion (Geffin and McCarthy, 2013).

Further complicating glial responses during progressive viral infection are the interactions between lymphocytes and monocytes that traffic into the nervous system during viral infection (Dou et al., 2006; Epstein and Gendelman, 1993; Gonzalez-Scarano and Martin-Garcia, 2005; Gorantla et al., 2010). Thus, a disease model that accurately replicates nervous system viral infection must contain both human immune and glial cell elements (Gorantla et al., 2012). This can be achieved by reconstitution of the mouse brain with human neural progenitor cells (NPC). Transplantable NPC readily produce neuronal and glial lineage cells in the rodent brain (Nunes et al., 2003; Uchida et al., 2000). The mice contain human hematopoietic stem progenitor cells (HSPC) readily generating lymphocyte and monocyte-macrophages susceptible to HIV-1 infection. Human glial (astrocytes and oligodendrocytes) evolve from intraventricular NPC transplantation at birth (Han et al., 2013; Windrem et al., 2014). Dual blood and brain humanized mice when infected with a macrophage-tropic CCR5 HIV-1_ADA_ can be examined for resultant effects on glial homeostasis. Transcriptional evaluations of two brain regions, the hippocampus and corpus callosum, demonstrated glial disease footprints in affecting antiviral human-specific immune responses. The importance of this model is underscored by specific glial transcriptional signatures that closely mirror the human disease (Gelman et al., 2012; Sanfilippo et al., 2017; Sanna et al., 2017).

## Results

### Human NPC engraftment

Mice were reconstituted with human immunocytes and glial cells. Newborn irradiated NSG mice were transplanted intraventricularly with human neuronal progenitor cells (NPC) and intrahepatically with cluster of differentiation 34 (CD34)^+^ HSPC from the same donor. NPC were isolated from fetal human brains to generate neurospheres, from which single cell suspension was prepared and transplanted at 10^5^ cells/mouse (experiments 1 and 2, n = 19) or 0.5×10^5^ (experiments 3 and 4, n=26). The fate and distribution of the human NPCs in mouse brain were then evaluated at 6 months of age. To analyze distribution of human glial cells paraffin embedded and formalin fixed tissue was sectioned along the sagittal plane from midline to the anatomical end of the corpus callosum (CC,∼ 2.5 mm according to adult mouse brain atlas) from left and right hemispheres. In two experiments (n = 19) the distribution of glial cells was observed. Human glial cells were found in equal measure across both hemispheres on sagittal sections with distributions in the periventricular area (PV), CC, ventral striatum (STRv), cerebral cortex (CCx), midbrain (MB), cerebellum (CB) and hippocampus (Hip) areas. Human cells were present in olfactory bulb (OB), lateral olphactory tract, anterior commissure, caudoputamen, midbrain, pons and medulla (Fig. 1B, C, D, Table 1). A large proportion of the human cells in mouse brains were GFAP positive with typical astrocyte morphology based on size and process. To distinguish human and mouse astrocytes the staining with non-species-specific (rb Dako/Z0034) and human-specific (ms STEM123) antibodies was done. Human astrocytes commonly replaced their murine counterparts with nearly 70–80% occupancy in PV, CC. (Fig. 1B, indicated by star). There was limited occupancy of human astrocytes in the frontal cortex (FC) and CB in 1/3 of analyzed mice (Fig. 1E). Surprisingly, nearly 40% of human nuclei protein (hN, MAB4383) positive cells in the CC and 20-30% in Hip and STR were Olig2^+^ (rb 1538-Olig2), an oligodendrocyte/progenitor transcriptional factor (Fig. 1D, F). The similar pattern (PV, CC, Hip, SRTv, MB) with less density of distribution of human cells were observed in two next experiments. To confirm glial lineage potential of the NPC we investigated their fate in short-term cultures. Mixtures of neural lineage cells arising from parental progenitors included GFAP/microtubule-associated protein (MAP) glial progenitors, nestin/GFAP neural progenitors (with radial glia characteristics), glutamic acid decarboxylase 67 (GAD67)-negative and class III β-tubulin (Tuj-1)^+^ neuronal lineage cells (Fig. S1). These data indicated that cells derived from neurospheres retained multi-lineage competence. Expanded human glial cells were integrated in mouse brain tissue without activation of human microglial cells (Fig. 2). We concluded that generated by neurosphere technique NPC transplanted at birth in brain lateral ventricle able efficiently repopulate NSG mouse brain with some variability, differentiate in astrocytes and retain oligodendrocyte development potential.

**Table 1.**
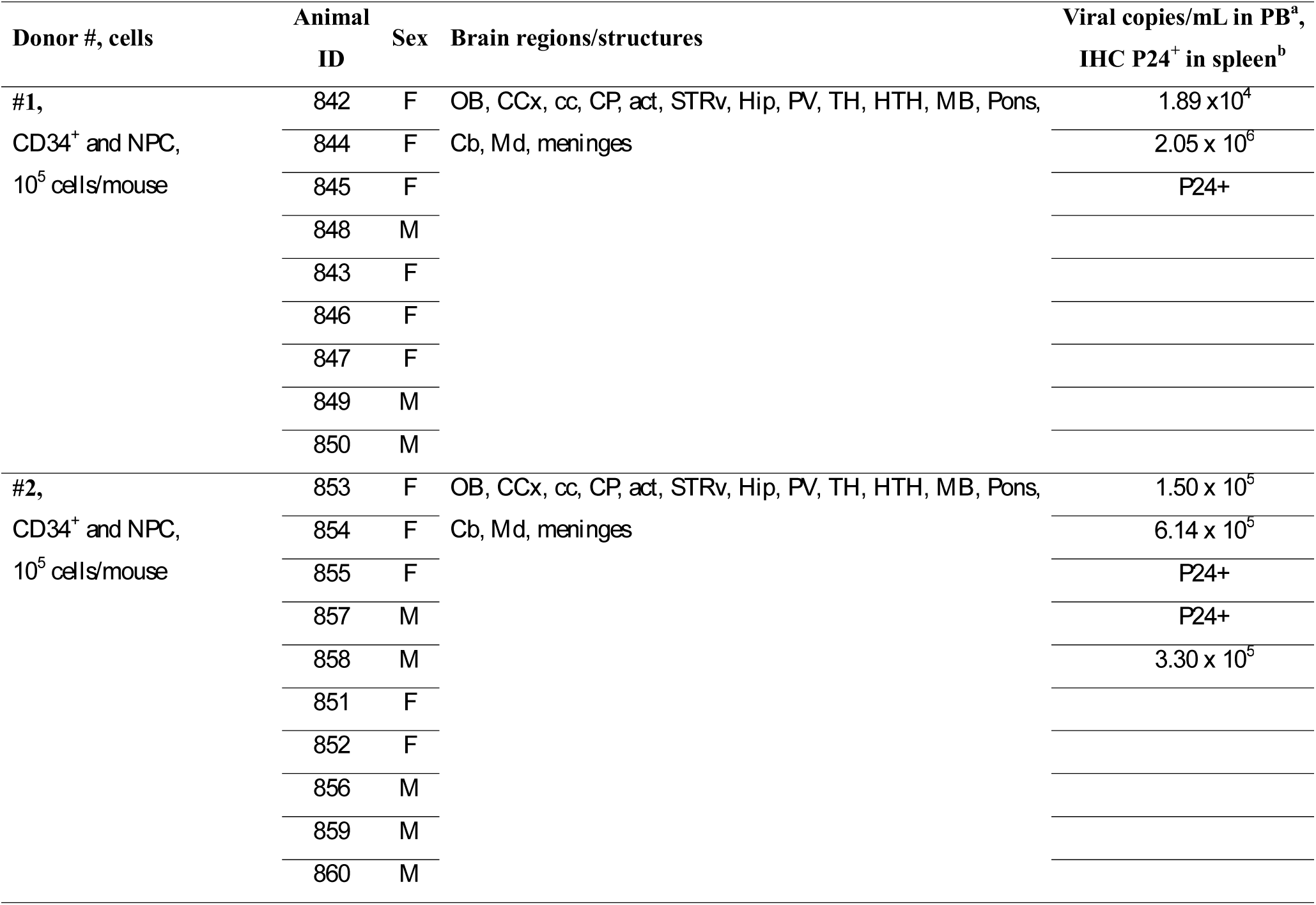
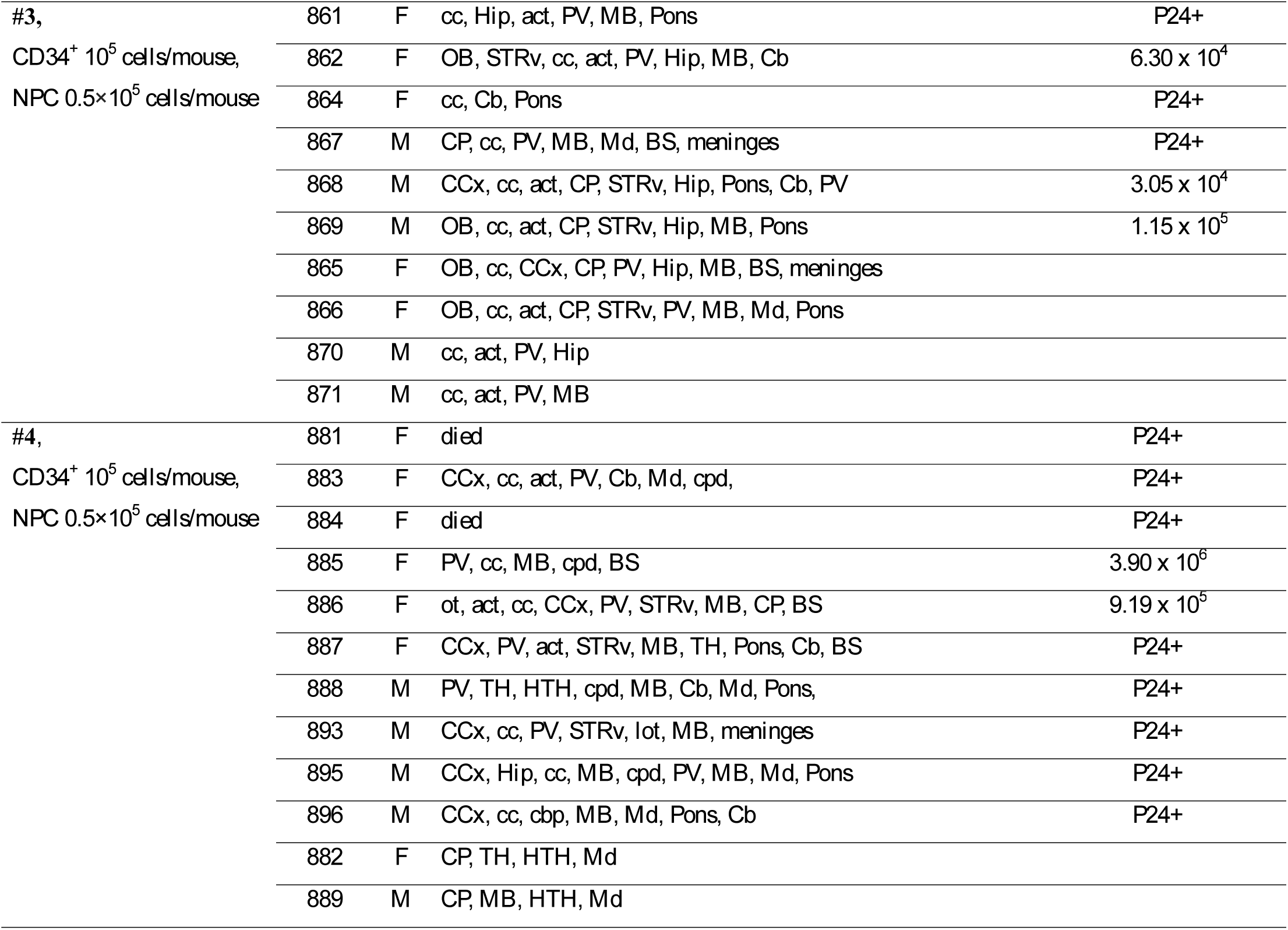
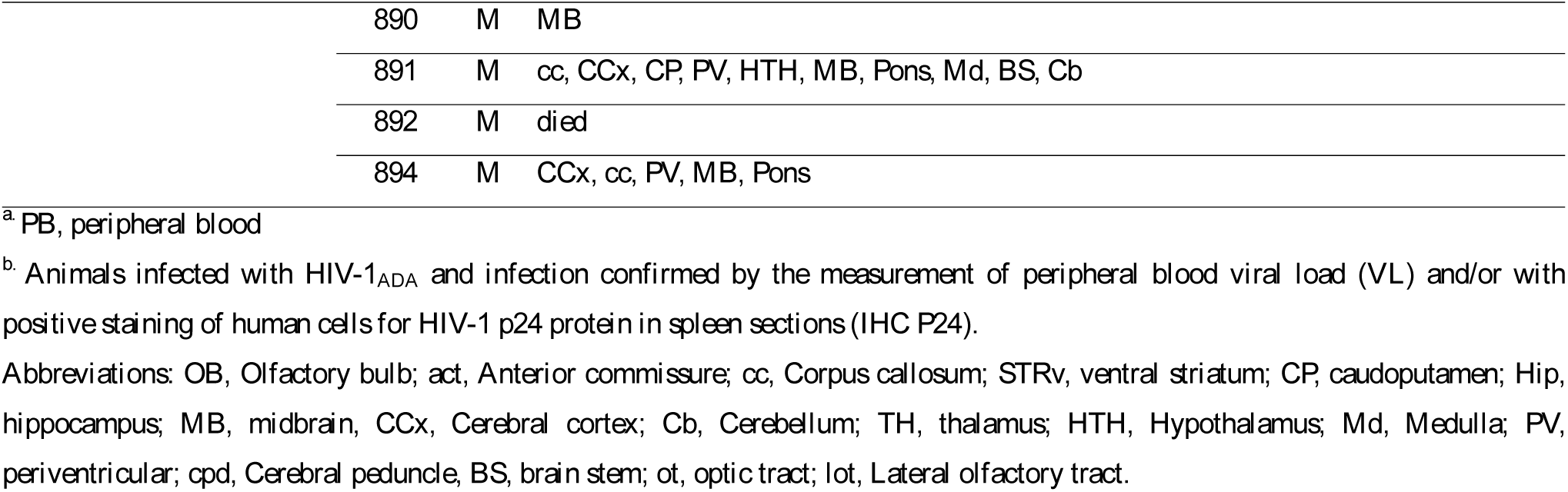
Human glial cell distribution pattern on sagittal plane.

**Figure 1.**
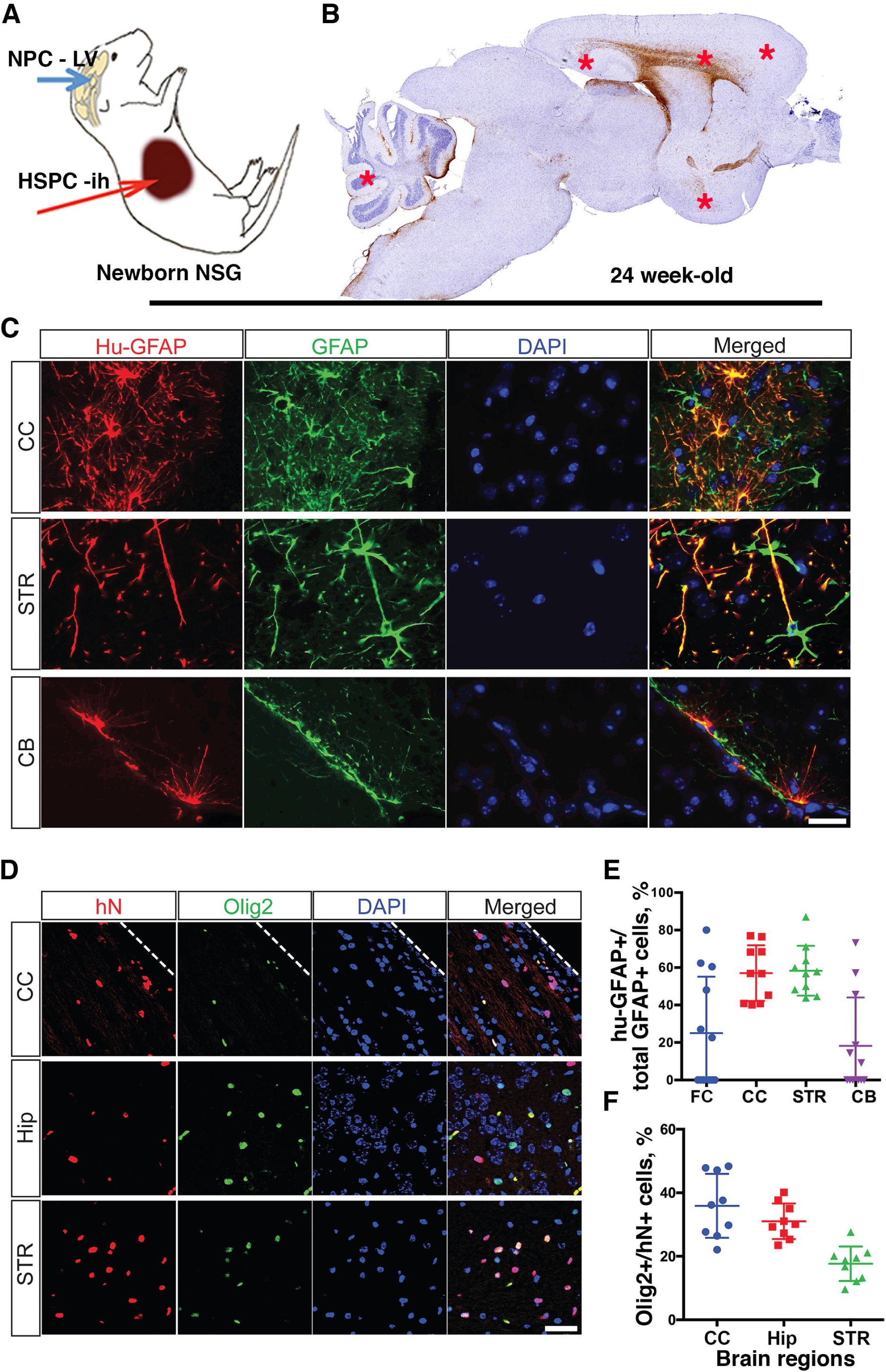
Glial humanization of the mouse brain. ***A,*** Scheme of the transplantation of human blood and brain progenitors at birth and the period of observation. ***B,*** Representative view under a 1× objective lens of 5-µm thick paraffin-embedded sagittal section stained for human-specific GFAP (brown). ***C,*** Human astrocytes specific antibodies (hu-GFAP, red) and non-species-specific total glial fibrillary acidic protein (tGFAP; green) staining was performed in different brain regions. Representative images are shown. Original images were collected by a Leica DM6 system (Leica Microsystems, Wetzlar, Germany) with 1000× magnification. Nuclei were stained with 4’,6-diamidino-2-phenylindole (DAPI; blue). Bar = 20 µm. ***D,*** Oligodendrocyte specific nuclear staining (Olig2; green) and total human-specific nuclear (hN, red) staining was done in different brain regions and representative images are shown. The original magnification was 400×. Bar = 50 µm. Dotted line on (**B**) demarcates the corpus callosum (CC) from the frontal cortex (FC). ***E, F,*** Quantitative data showing the number of human glial cells from one of four experiments with ten animals. The average percentage of human cells was assessed on 2–4 representative sagittal sections from each mouse for a selected field of view of the same brain region under 400× magnification (indicated by star). ***E***, Distribution and proportion of hu-GFAP^+^ astrocytes among total number of GFAP^+^ cells. ***F***, Distribution and proportion of hN^+^ cells positive for Olig2. Each symbol represents an individual mouse and lines indicate mean and standard error of the mean (SEM). Animals were analyzed at 6 months of age and reconstitution characteristics is shown in Table 1.

**Figure 2.**
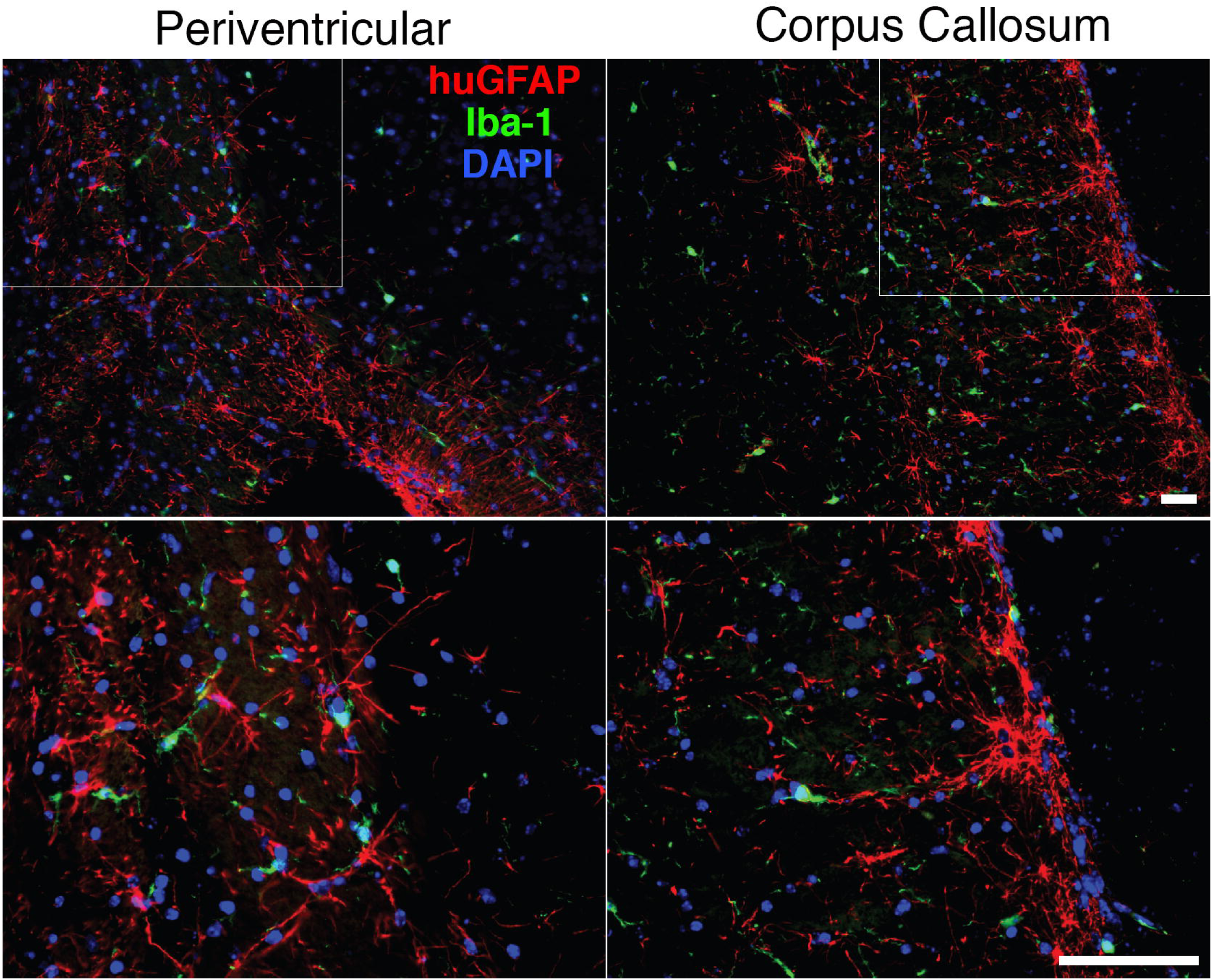
Interaction of human and mouse glial cells. Sagittal section of paraffin-embedded brain tissues was stained for human astrocytes (hu-GFAP, red) and mouse microglial cells (Iba-1, green) identification. Original images in periventricular region and corpus callosum were collected by a Leica DM6 system (Leica Microsystems, Wetzlar, Germany) with 100× magnification. Selected fields of view (square) were captured with 400× magnification. Bar = 100 µm. Mouse microglial cells are in contact with human astrocytes. There were no morphological signs of activation, formation of microglial nodules or active phagocytosis of human cells.

### Efficient peripheral hemato-lymphoid humanization of NSG mice

A human peripheral hemato-lymphoid system was established in the NSG mice (Fig. 3). Flow cytometric analyses of peripheral blood of humanized mice at 24 weeks of age showed that among human CD45^+^ cells, B (CD19^+^) and T lymphocytes (CD3^+^) were present (Fig. 3A). The CD14^+^ monocyte population in peripheral blood were also present (Fig. 3B). Human hemato-lymphoid reconstitution was similar to previously published (Arainga et al., 2016; Dash et al., 2011; Gorantla et al., 2010). In the brain, human peripheral blood leukocytes infiltrated the meningeal and perivascular spaces, parenchyma. The majority of HLA-DR cells were located in the meningeal space, with fewer numbers in the perivascular space and brain parenchyma (Fig. 3C and D). As we published previously (Gorantla et al., 2010), among these, the greater part were CD163^+^ macrophages, and some showed branching appearances. Less than 20% of HLA-DR cells were CD4^+^, and 9% were CD8^+^ T lymphocytes (Fig. 3C and E). The data were collected in animals with similar levels of human cells in peripheral blood (n = 7).

**Figure 3.**
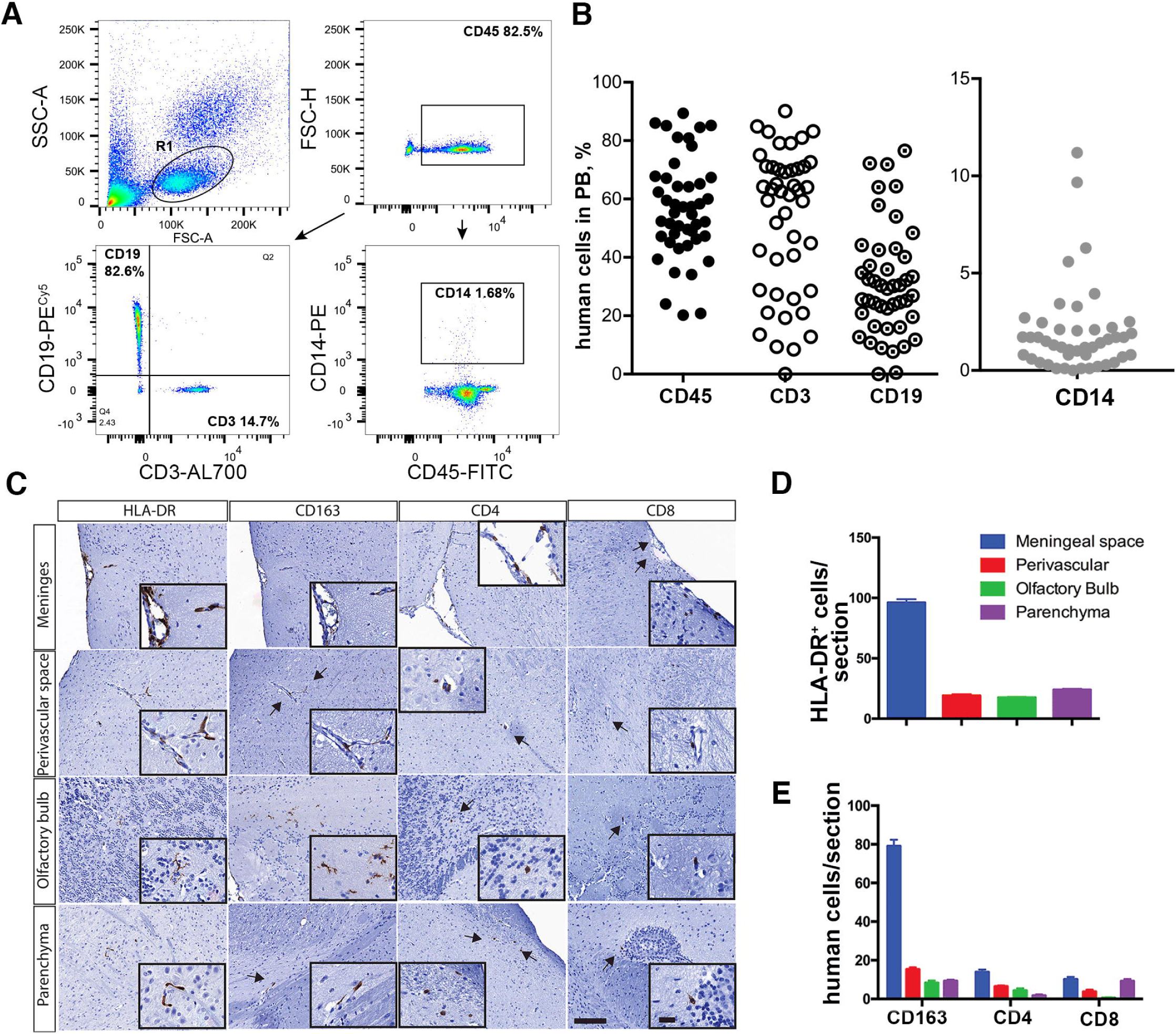
Human immune cell repopulation of mouse blood and brain. ***A,*** Fluorescence-activated cell sorting (FACS) analysis of peripheral blood. Representative plots of human cluster of differentiation 45 (CD45)^+^ cells and human CD3, CD19, and CD14 cells gated from the CD45 lymphocyte population. ***B,*** Percentage of human cells in peripheral blood of all dual reconstituted mice used in the studies. Each symbol represents an individual mouse. ***C,*** Representative images of the distribution of human immune cells in brain are shown. Paraffin embedded 5-µm thick sections were stained for human-specific markers: human leukocyte antigen-antigen D related (HLA-DR), CD163, CD4, and CD8. Representative images from different brain regions are shown. Images were captured by a Nuance multiplex system (CRi, Wobum, MA) with an original magnification of 200×. Insets represent cells with a magnification of 1000×. Bars = 100 µm and 20 µm (inset). ***D*** and ***E***, Quantification of human cells positive for HLA-DR, CD163, CD8, and CD4 in selected regions. Numbers represent frequencies of positive cells per section; 7 mice with similar levels of lymphoid reconstitution were analyzed.

### HIV-1 infection of lymphoid and brain tissues in dual humanized mice

At 24 weeks of age, mice were divided into two groups. One group was infected intraperitoneally with HIV-1_ADA_ at 10^4^ infectious particles in a volume of 100 µl. We did not observe a significant decline in human CD4^+^ cells at the time of infection. At ∼5 weeks flow cytometric analyses showed ∼ 20% drop in circulating CD4^+^ T cell numbers among the CD3^+^ cell population (p = 0.023) in selected animals with similar levels of human cells in peripheral blood (n = 6/group). An increase in the CD8^+^ T lymphocytes in proportion to CD4^+^ T cells was observed (Fig. 4A and B). After infection, HIV-1 p24^+^ stained cells were present in spleen (Fig. 4C) and mice showed a median of 1.6 × 10^5^ viral RNA copies/ml in blood (Fig. 4D). The brain of infected animals (frontal cortex and striatum) were analyzed for the presence of viral RNA (Fig. 4E). The levels of viral RNA expression in the brain tissue remain low (< 50 copies/µg of RNA) compare to systemic infection. HIV-1infected human cells were observed in meninges (Fig. 4 F). HIV-1 infection influenced immunocyte infiltration in the brain and glial reconstitution. Numbers of human immunocytes were analyzed for animals with comparable levels of peripheral blood leukocyte repopulation by counting HLA-DR^+^ and CD163^+^ cells/sagittal section in HIV-1-infected (n = 5), uninfected (n = 6) dual transplanted. A statistically significant two-fold increase in the number of brain infiltrating immunocytes following HIV-1 infection was observed in meninges compared to uninfected dual reconstituted mice 218.2 ± 8.6 versus 112.0 ± 13.3 cells/section (p = 0.0002). In the brain parenchyma activated human HLA-DR^+^ cells were 29.8 ± 2.8 and 17.0 ± 3.2 cells/section in infected and uninfected mice, respectively (p = 0.017). The majority of human immunocytes were CD163^+^ macrophages (Fig. 4G). In the spleen HIV-1p24 cells were present (Fig. 4C), and proportion of stained cells for the HIV-1 Nef and p24 proteins in spleen were similar (not shown). To analyze changes in glial cells the animals from two experiments with similar and broad distribution of human glial cells were selected. These sections were stained for human-specific GFAP, counterstained with hematoxylin and then subjected to unbiased digital image analysis (DEFINIENS Tissue Studio®, Cambridge, MA, USA) to assess human glial cell repopulation in 5-µm sections. The total number of nucleated cells/sagittal section remained consistent across the animal groups 107,061 ± 5,147 and 110,908 ± 6,483 cells/section (Fig. 4H). In comparison to uninfected controls (n = 18), HIV-1-infected mice (n = 19) had a lower proportion of human astrocytes/sagittal section 20.7 ± 1.5 % versus 15.9 ± 1.4 % of human-specific GFAP^+^ cells, respectively (p = 0.022). Serial sections also were stained for hu-GFAP, HIV-1p24 and Nef. No astrocytes were identified stained with p24 and Nef viral proteins (data not shown). Systemic HIV-1 infection of dual humanized animals increased influx of human macrophages and affect number of hu-GFAP expressing cells.

**Figure 4.**
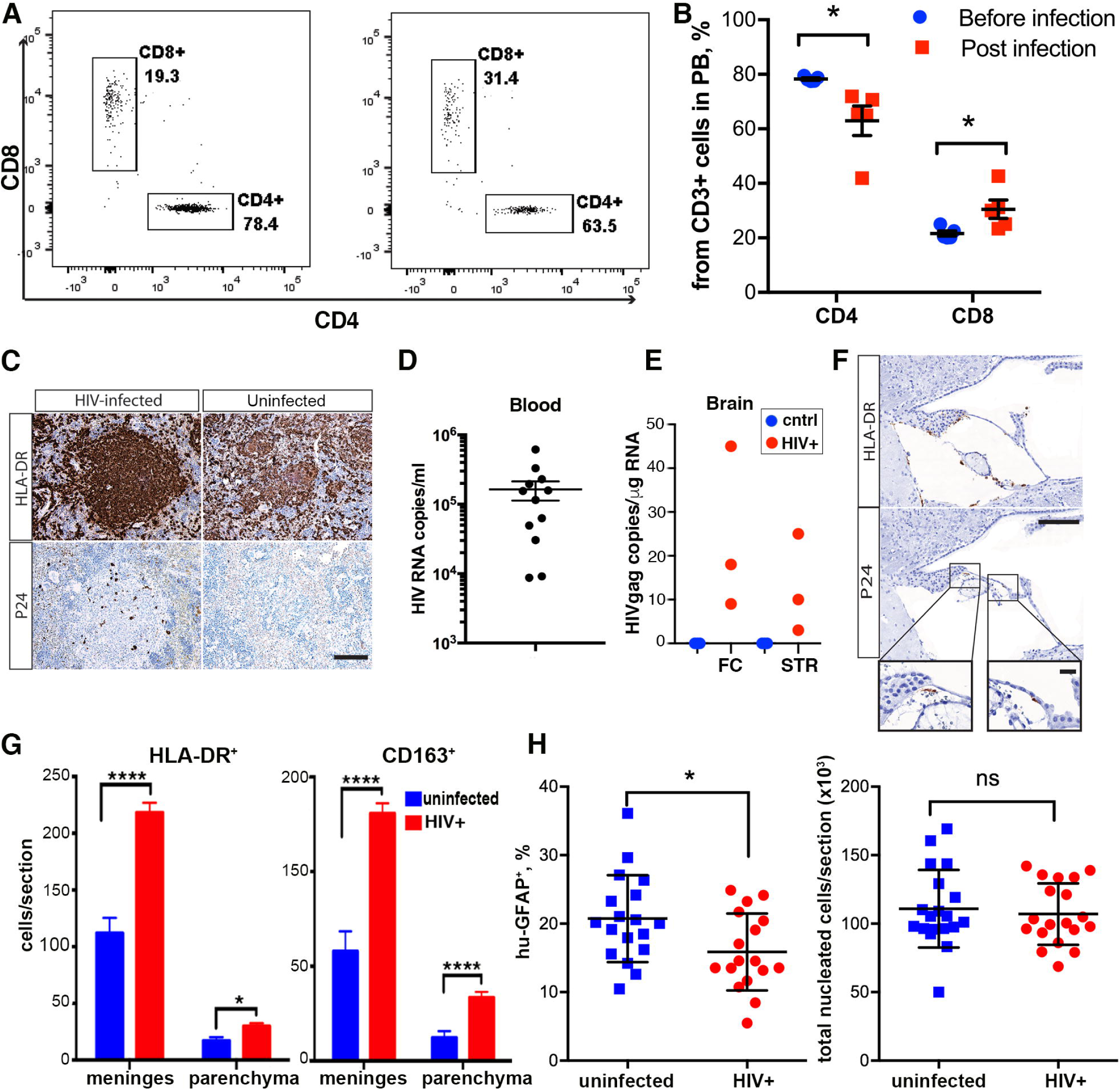
HIV-1 infection affects human cell distributions in dual blood and brain humanized mice. ***A***, Examples of representative plots showing CD4^+^ cell decline in peripheral blood of HIV-1-infected mice, with a relative increase in the CD8^+^ cell compartment. ***B,*** Quantification of human CD4 and CD8 T cells in the peripheral blood of infected (n = 6) and control (n = 6) mice. Multiple t-tests using the Holm-Sidak method for multiple comparisons correction was applied to examine significant differences between groups. Bars = means ± standard error of the mean (SEM); *Adjusted P = 0.045. ***C***, Immunohistology of spleen sections shows HLA-DR positive cells and the presence of HIV-1-infected cells stained for HIV-1 p24. Images were captured by a Nuance multiplex system (CRi, Wobum, MA) with an original magnification of 200×. Bar = 100 µm. ***D,*** Peripheral viral load determined by a COBAS Amplicor System v1.5 (Roche Molecular Diagnostics, Pleasanton, CA, USA). Each symbol represents an individual infected mouse from all experiments. ***E***, Viral RNA levels were determined by semi-nested RT-PCR in the HIV^+^ group (n = 3, red symbol) and the control group (n = 3, blue symbol) in two brain regions, frontal cortex (FC) and striatum (STR). ***F***, Immunohistology of brain sections showing the presence of activated HLA-DR^+^ cells and HIV-1p24^+^ infected cells on adjacent serial section. Representative images from the meningeal space containing vascular vessels are shown. Images were captured by a Nuance multiplex system with an original magnification of 200×. Insets represent cells with a magnification of 1000×. Bars = 100 µm and 20 µm (inset). ***G***, Quantification of infiltrating HLA-DR^+^ and CD163^+^ cells in the brain regions of HIV-1-infected (n = 5) and uninfected dual reconstituted mice (n = 6). Two-way analysis of variance (ANOVA) using Sidak’s multiple comparisons correction was used to examine significant differences between groups. Bars = means ± SEM; *P = 0.017, ****P < 0.0001. ***H***, Quantification of human glial fibrillary acidic protein (GFAP)^+^ and all nucleated cells on representative sagittal brain sections of HIV-1-infected (n = 18) and control (n = 19) mice. Unpaired t-tests were used to examine significant differences between groups. Each point is individual animals averaged counts from two sections presented, means ± SEM are shown; *P = 0.022, ns = no significance.

### Transcriptional changes in the humanized mouse brain induced by HIV-1 infection

To assess the pathobiological outcomes of viral infection following peripheral immune and brain humanization, we applied deep sequencing to evaluate the transcriptional profiles seen as a consequence of virus-immune cells-glia crosstalk. We selected three uninfected and three infected animals with similar patterns of human immune cell repopulation, similar pattern of astrocyte distribution and density, and similar peripheral viral loads (for infected mice) for study (Fig. 5A). Three equivalent aged unmanipulated NSG mice served as controls. Hip and CC repopulated by human glial cells were dissected for sequencing (Fig. 5B). All sequenced reads (∼100 bp length) were aligned to the mouse database University of California Santa Cruz Genome Browser database (GRCm38/mm10). The experiment was designed to search for differentially expressed mouse genes as a result of human cell engraftment and HIV-1-infection (Breschi et al., 2017). Subsequently, the reads unmapped to the mouse genome were re-aligned to the human genome (GRCh38/hg19) to find human-specific transcriptional changes. The overall mapped mm10 reads average ratio was 92% for all 9 Hip samples and 94% for all 9 CC samples. For the HIV-1-infected and uninfected humanized mice, the unmapped to mouse database reads were run against its human genome. The average ratio for mapped reads was 32% and 26% for the Hip and CC, respectively. Analyses using the mouse database identified 365 differentially expressed transcripts in the Hip and 93 in the CC as a result of the engraftment of human glial cells. When compared to humanized mouse transcripts, few mouse transcripts were differentially expressed as a result of HIV-1 infection in Hip only (Fig. 5E). When the reads unmapped to the mouse genome were realigned to the human reference genome, 115 and 58 human transcripts were found differentially expressed in the Hip and CC, respectively, following infection (Fig. 5F and G). The expression (FPKM) of human lineage-specific genes CD163, CD4 (macrophages and lymphocytes) were significantly lower compared to human glial transcription factors OLIG1, OLIG2 (progenitors and oligodendrocyte commitment), GFAP and Glutamate-Ammonia Ligase (glutamine synthase family), genes expressed by astrocytes. HIV-1 infection reduced expression of human GFAP (FPKM log_2_ changes −0.55, p = 0.033). Expression of mouse cell lineage-specific markers was not statistically significantly affected (data not shown). Region-specific alterations of mouse transcriptome by engrafted human glial cells we summarized in Supplemental Table 1, Fig. S2 and S3). Glial humanization of mouse brain induced regions-specific transcriptional changes associated with multiple biological processes and is of interest beyond this report (Goldman et al., 2015).

**Figure 5.**
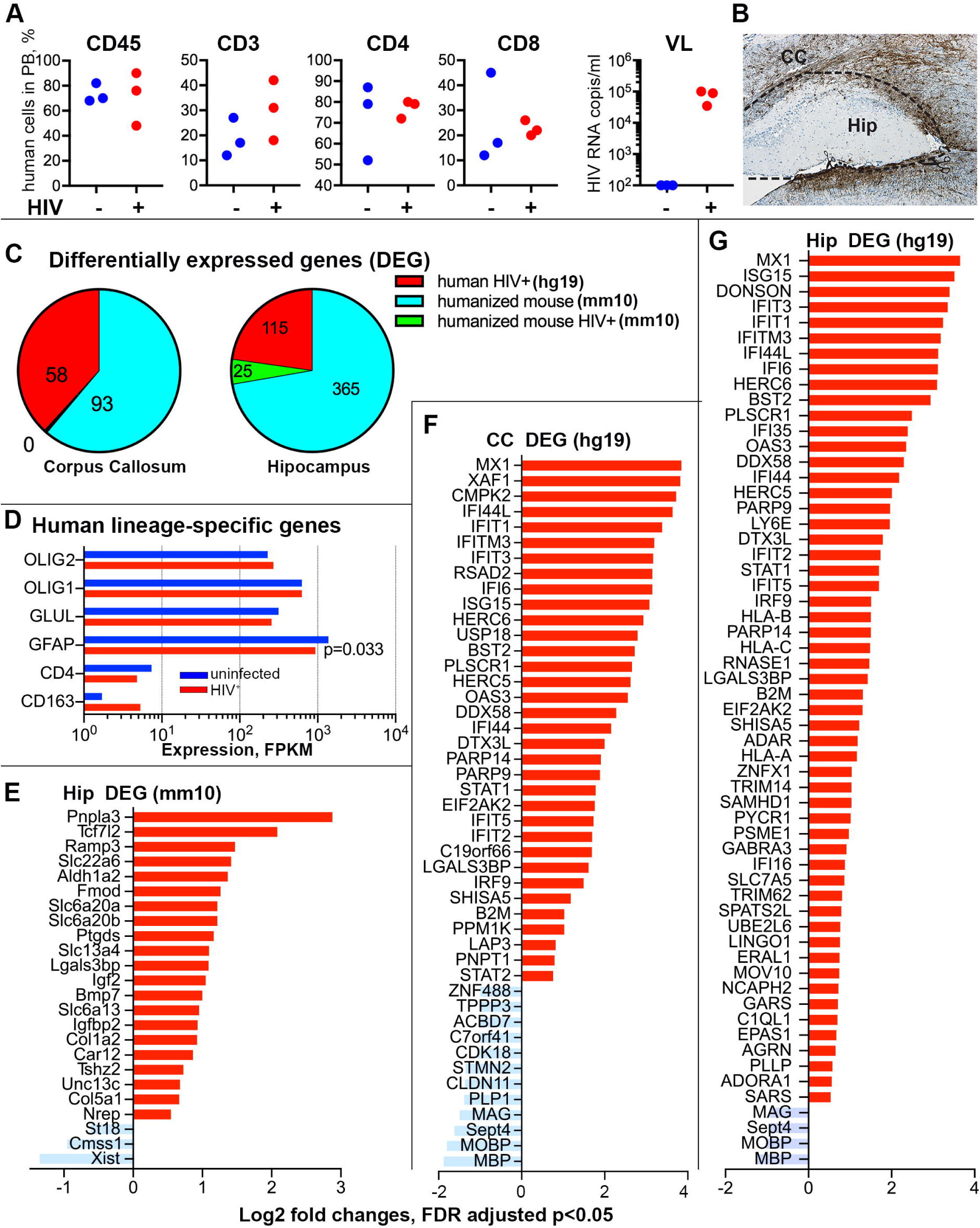
HIV-1 infection induces transcriptional alterations in brains of dual humanized mice. ***A***, Characteristics of peripheral reconstitution and the levels of HIV-1 infection of mice selected for RNA-sequence analysis (n = 3 per group). Percentage of human CD45, CD3, CD4, and CD8 cells, as well as HIV-1 viral load (VL) in the peripheral blood of HIV-1-infected dual reconstituted mice (HIV^+^, red symbols) and control dual reconstituted mice (control, blue symbols) are shown. Three unmanipulated NSG of the same age were used as a control and were negative for all these parameters. ***B***, Representative image of human glial cell reconstitution and scheme of CC and Hip microdissection. ***C***, Identification of differentially expressed genes. Sequenced reads were mapped to the mouse database (mm10) and comparisons were conducted: 1) between non-manipulated NSG mice and dual reconstituted NSG mice to find differences induced by humanization (humanized mouse mm10, blue color); 2) between uninfected and HIV-1 infected dual reconstituted mice to observe differences of mouse transcriptome induced by infection (humanized mouse mm10, green) and in CC such differences were not identified. Reads unmapped to the mm10 database was realigned to the human database (hg19) and comparisons were conducted between uninfected and HIV-1-infected dual reconstituted mice (human HIV^+^ hg19, red color). ***D***, Confirmation of the expression of human cell lineage-specific genes for Hip (hg19) and effects of infection on human GFAP downregulation were statistically significant in t-test comparison, P = 0.033, but did not reach significance with FDR adjustment. None of mouse cell lineage-specific genes expression was affected by HIV-1 infection. ***E***, Up- and downregulated mouse genes in the Hip of HIV-1-infected dual reconstituted mice compared to uninfected mice (mm10). ***F***, ***G,*** Up- and downregulated human genes in the CC and Hip of HIV-1-infected dual reconstituted mice compared to uninfected mice (hg19). ***E, F, G***, Genes with 1.5 fold changes with FDR adjusted p < 0.05 were graphed.

**Figure 6.**
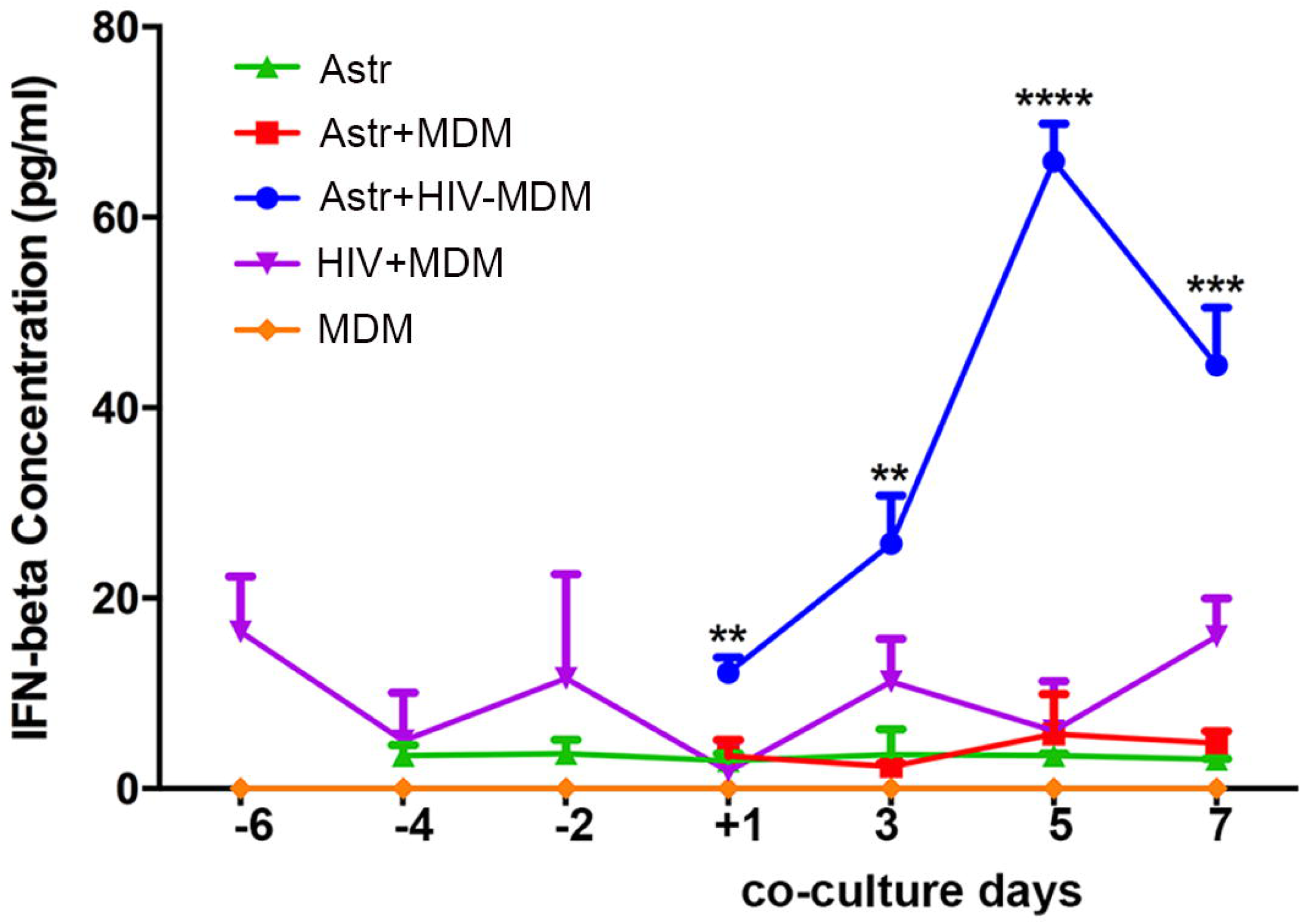
Astrocyte production of IFN-β. Levels of human IFN-β were measured by enzyme-linked immunosorbent assay (ELISA) in the supernatants collected from astrocytes (green), a co-culture of astrocytes and control monocyte-derived macrophages (astrocytes and MDM, red), co-culture of astrocytes and HIV-infected macrophages (astrocytes and HIV-1 infected MDM, blue), HIV-1infected and control MDM purple and orange, respectively). X-axis indicates the time points before and after co-culture (24 hours, 3, 5, and 7 days). Data are expressed as the mean of triplicate technical replicates from one from three independent experiments (each done with distinct donors). Student t-tests were performed to compare Astrocytes with and without HIV-1 infected MDM (red and blue) for each time point (2 and 4 days). Bars = means ± standard error of the mean (SEM); **p = 0.0029, ***P = 0.0004, and ****p < 0.0001.

### Alterations in the mouse transcriptomes following viral infection

HIV-1-induced transcriptional changes analyzed with alignment to mouse genome identified 25 DEG (Fig. 5E) and according to GeneCards® integrative database identified Gene Ontology (GO) biologic processes gene sets could be associated with metabolic processes, blood vessel development and extracellular matrix re-organization (*ALDH1A2, TCF7L2, COL1A2, COL5A1, FMOD, BMP7*) as well as neurotransmitters, sugars, organic acids, anion transmembrane transport (*SLC6A13, SLC6A20, SLC22A6, SLC13A4*), and cellular responses to hormone stimulus (*IGFBP2, RAMP3*). Few downregulated genes were observed (*ST18, XIST*) and expression levels of these genes could be found in supplemental data (Supplemental Table S2). The changes in the neurotransmitters expression in Hip of infected humanized mouse brains could affect cognition (Zhou and Danbolt, 2013) and need further evaluation. For example, downregulated by humanization expression of *IGFBP2* was up-regulated by HIV-1 infection and associated with inflammatory responses in HIV-1 infected patients (Helle et al., 2001; Suh et al., 2015). Another example is downregulation of *XIST* initially upregulated by glial humanization and related to the control of gene expression (Groen and Morris, 2013). These changes do not reflect ongoing viral infection.

### Species-specific alterations in the human transcriptomes following viral infection

The major observation is species-specific transcriptional changes in dual humanized HIV-1 infected mice brains. In contrast to mouse transcriptome, alignment to human genome reveals overlapping pattern of upregulated and downregulated genes in Hip and CC (Fig. 5F and G, Fig. S3, Supplemental Table S2). The upregulated genes were strongly associated with type 1 and 2 IFN responses and specific signaling pathways. These were linked to host viral defense responses (*MX1, IFIT* families, *PLSCR1, EIF2AK2, IFI16, ADAR, OAS3, MX1, BST2, RSAD2*), RNA catabolism (*CMPK2, PNPT1*), ISG15-protein conjugation, regulation of type 1 IFN production (*HERC6, ISG15, DXT3L, DDX58, PPAR9, PPM1K*) and cytokine-mediated signaling. The spectrum of Hip upregulated genes overlapped with that of the CC. In addition, upregulated genes associated with norepinephrine metabolic processes (*EPAS1, LY6E*) and those related to neural and glial differentiation (*PARP10, GARS, AGRN, SOX10, LINGO1*) were found. In the CC and Hip, the identified downregulated genes were linked to myelination and axon ensheathment (*MBP, MOBP, MAG, PLP1, CLDN11*), neural progenitor differentiation (*STMN2, ZNF488, IGFBP5, BIN1, APLP1*), glial structural integrity, lipid metabolism, ion transportation and cell cycle regulation (Fig. S3, Supplemental Table S2). Identified transcriptional changes show ongoing human glial antiviral response to HIV-1 infection.

### Human astrocytes produce type 1 IFN during infected macrophage-astrocyte co-cultivation

The upregulated expression of genes related to type 1 IFN responses in the chimeric murine brains following HIV-1 infection need investigation of the possible source of interferon. Though almost all nucleated cells can produce type 1 IFN, plasmacytoid dendritic cells are the most potent IFN producers (Steinman, 1991). However, prior studies have demonstrated that astrocytes abortively infected by neurotropic viruses are also a potent source of IFN beta (IFN-β) in mice (Pfefferkorn et al., 2016). To identify the source of type 1 IFN in response to HIV-1 infection in dual humanized mice, we used transwell systems to co-culture human fetal astrocytes and HIV-1-infected monocyte-derived macrophages (MDM). After 3 days of co-culture with HIV-1-infected MDM and astrocytes, astrocytes were found to produce IFN-β. The levels of IFN-β reached a peak at 5 days. Three independent donors were used; a representative experiment is shown in 6. The concentration of IFN-β produced by HIV-infected MDM alone remained at relatively low levels (approximately 15 pg/ml, detection limit). The concentration of IFN-β in control astrocytes and MDM were below detectable throughout the experimental observation period.

### RT-PCR confirmation of canonical interferon stimulated human genes expression in humanized mice brain tissues

To confirm the altered expression and the species-specificity of the IFN signaling pathway related transcripts, we applied real-time PCR assays to reexamine the selected genes found affected in RNA-sequencing using human and mouse specific primers and probes. As critical elements of the upstream of type 1 IFN response pathway, *STAT1, STAT2* and *IRF9* were found 3.72, 1.69, and 3.64 times (p = 0.0003, 0.039, 0.00002) upregulated in the CC of HIV-1-infected dual reconstituted mice, respectively, compared to that of control dual reconstituted mice. Upregulation of these three genes was found in the Hip of HIV-infected humanized mice as well (Fig. 7, p = 0.004, 0.020, 0.010). The down-stream IFN inducible genes such as *MX1, ISG15, IFI6* and *CMPK2* were also found significantly upregulated in both CC (12.09, 11.69, 9.41 and 13.69 times, respectively, p< 0.000001) and Hip (19.90, 15.90, 9.17 and 21.25 times, respectively, p< 0.000001) of HIV-1 infected humanized mice compared to the control animals. We confirmed species-specific upregulation of IRF9/STAT1/STAT2 interferon stimulated response elements that mediate resistance to viruses (Cheon et al., 2013). Among the myelination and glial integrity associated genes, the expression of *MBP* was reduced 2.94 and 2.5 times (p = 0.0058, 0.0013) in the Hip and CC of HIV-1 infected mice, respectively. The expression of ZNF488 was only found significantly downregulated in the CC of HIV-1 infected group (p = 0.0129), but not in the Hip. Mouse MBP RT-PCR failed to demonstrate gene expression difference between the HIV-1 infected and control animal groups (Fig. 7).

**Figure 7.**
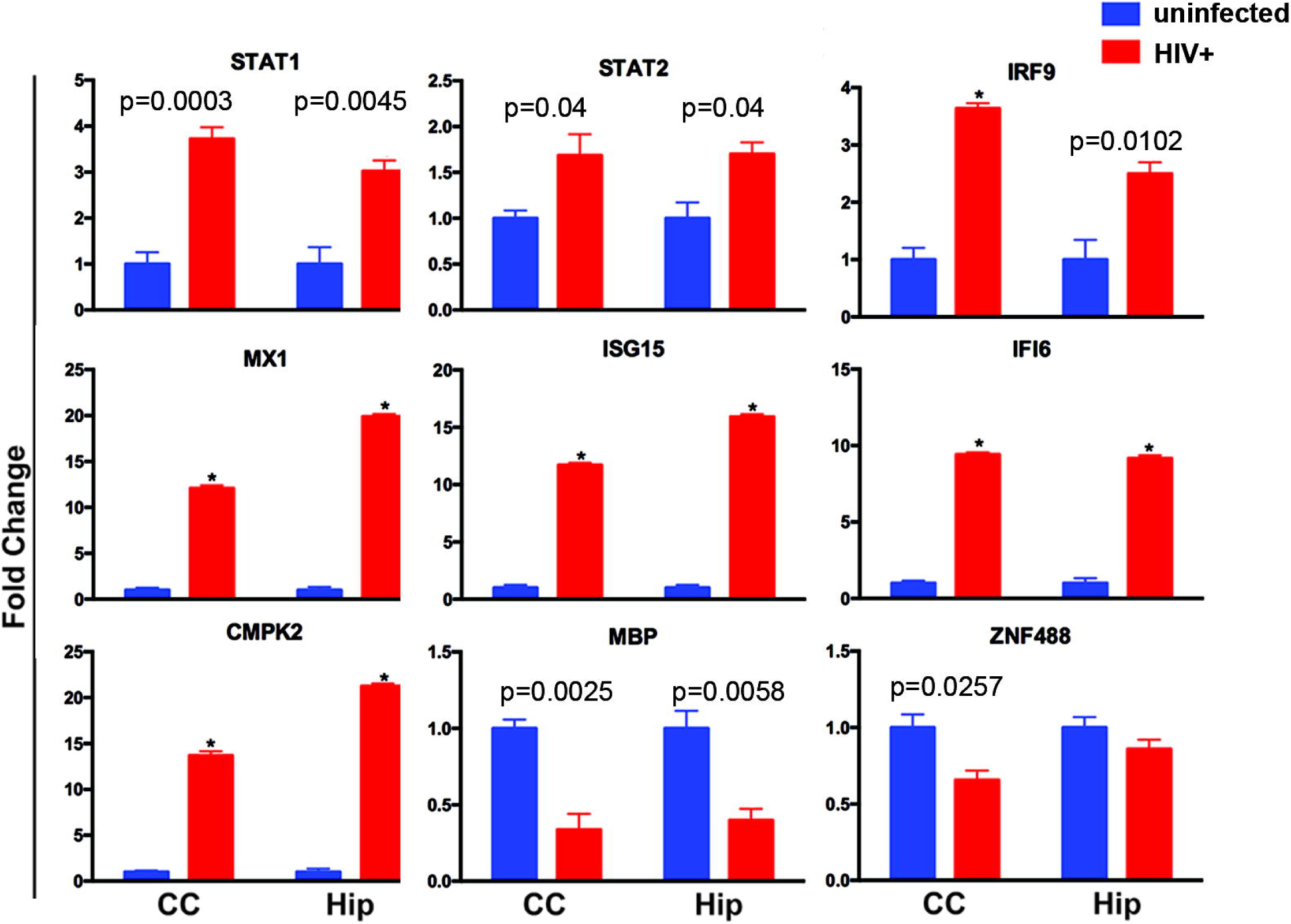
Real-time PCR confirmation of affected human genes in humanized mice brain tissues. Selected human gene expression profiles in the CC and Hip were confirmed by TaqMan® real-time PCR. The HIV-1 infected (n = 6) and control group (n = 5) are illustrated. The fold change in gene expression of human *STAT1, STAT2, IRF9, MX1, ISG15, IFI6, CMPK2, MBP,* and *ZNF488* between the HIV-1 infected and the control groups were determined by using the 2^-ΔΔ*C*^T method (A-I), where ΔCt = Ct_target_ – Ct_GAPDH_ and Δ(ΔCt) =ΔCt_HIV_ – ΔCt_control_. Multiple t-tests using Holm-Sidak method for multiple comparisons correction was applied to examine significant group differences. Bars = means ± standard error of the mean (SEM); * - FDR adjusted p < 0.00001.

## Discussion

A mouse model reflecting HIV-1 infection of the nervous system was created that contains both human immune and glial cells in lymphoid organs and brain, respectively. This model represents the combination of NPC transplantation to generate human glia (Nunes et al., 2003; Uchida et al., 2000) and CD34^+^ HSPC reconstitution to forge a human immune system (Ito et al., 2002). This model builds on previous work demonstrating that transplantation of human neurosphere-derived NPC can occur in a living mouse with functional human-derived neurons and glia (Tamaki et al., 2002; Uchida et al., 2000). Early transplantation proved essential to optimize migratory responses of human glia in the mouse brain to facilitate replacement of their murine counterparts through brain subregions (Han et al., 2013; Windrem et al., 2014). We successfully show that human NPC when transplanted at an early postnatal age demonstrate broad glial distribution through the brain. By 6 months of age, considerable numbers of human astrocytes were seen to have repopulated multiple brain regions along whiter matter tracts. Moreover, we found considerable numbers of human Olig2 positive cells (Fig. 1) and signaling the emergence of significant oligodendrocyte differentiation Olig1 (Fig. 3D). The broad variability in glial cell distribution and density were associated, in part, with the number of transplanted cells, NPC gestation and collection times and their stem cell properties that are effected by the number of cell passages and the accuracy of intraventricular injections.

The distribution of human immunocytes in the brain vary and were comparable to what was observed in humanized mice created by CD34^+^ cell transplantation and subsequent HIV-1 infection (Gorantla et al., 2010; Honeycutt et al., 2016). Notably, the majority of human immunocytes present in the brain were observed in the meninges. These were predominantly CD163-positive human macrophages. High levels of systemic HIV-1 infection with infected meningeal and brain-infiltrating cells at lower numbers than what were seen in peripheral tissues were detected. Thus, the model reflects three principal features of human brain disease that includes a functional immune system with mobile brain penetrant monocyte-macrophages, sustained high levels of systemic viral infection and the presence of glial target cells that determine alterations in the brain microenvironment. Most importantly, each of the cell types were able to interact, one with the other, in a physiological manner. In contrast to a number of operative disease models, the effects of HIV-1 infection on human brain cells and human-specific immune changes that result from persistent systemic infection can now be fully evaluated.

We applied RNA-sequencing for dissection of species-specific changes to assess the relevance of the model to human disease. Indeed, our findings in the transcriptome of HIV-1-infected dual reconstituted mice corroborated the relevance of the animal model system. As the most abundant cell type in the brain, astrocytes react to injury or viral infection through cellular activation (Sabri et al., 2003). While HIV-1 infection of CD4^+^ T lymphocytes, perivascular macrophages and microglia are well recognized (Yadav and Collman, 2009), infection of astrocytes remains unresolved. The question that has remained could low level infection of astrocytes or mere exposure of astrocytes to infected cells or HIV-1 itself cause important physiological changes that alter the brain’s functional capacities? Indeed, in contrast to high levels of systemic infections, low levels of HIV-1 RNA were seen in analyzed tissue samples. While restriction of astrocyte infection can occur at the level of cell entry (Gorry et al., 2003), this in no way detracts from the notion that significant innate immune responses could still occur by the limited virus that is present and explain what was observed at transcriptional levels. Indeed, more than 30 human genes significantly upregulated in the CC and Hip was found to be closely related to innate and adaptive immunity, HIV host restriction factors, and IFN stimulated factors (Fig. 5, Fig. S3). These findings support the idea that low levels of virus can induce considerable affects in the brain’s innate antiviral immune responses. Noticeably, STAT1/STAT2 and IRF9 form a transcriptional complex upon activation of type 1 IFN receptors; this complex can subsequently translocate to the nucleus and induce the expression of a broad spectrum of IFN-stimulated genes that serve to contain infection (Schreiber, 2017). Among such IFN stimulated factors, *MX1, OAS3, RSAD2* (viperin), and all the *IFI* and *IFIT* family are involved in combating viruses at nearly every stage of the viral life cycle (Espert et al., 2005; Fricke et al., 2014; Laguette et al., 2011; Lu et al., 2011; Nasr et al., 2012; Neil et al., 2008; Okumura et al., 2006; Rivieccio et al., 2006; Spragg and Emerman, 2013). In the Hip, in addition to the upregulated genes introduced previously, class I HLA genes were found upregulated, which is yet another downstream effect of the type 1 IFN response.

The upregulation of the human specific IFN signaling related genes IRF9/STAT1/STAT2 were confirmed by real-time PCR assays using human specific probes. We also confirmed by RT-PCR upregulation of rare mRNA transcript identified by RNA sequencing for *CMPK2*, which could belong to infiltrating human macrophages as well as glial cells, and further affirm that the factors serve as a viral restriction factor (Lazear et al., 2013). However, no change in expression of the host restriction factors were reflected in the mouse transcriptional profile as result of HIV-1 infection. This confirms that HIV-1 can neither efficiently enter mouse cells due to the absence of specific receptors, nor trigger a pattern recognition response by murine cellular defense machinery after viral infection. We cannot exclude that extension of the observation time in the fact that few months of infection could be required to trigger mouse cell responses to HIV-1 and as such change the profile of human cell responses and damage.

This vigorous type 1 IFN signaling response further leads to the question of what cell source produces type 1 IFN in these dual reconstituted mouse brains. Due to the increased infiltration of macrophages into the brain during systemic HIV-1 infection in these mice, we tested these two-cell types interaction for type 1 IFN production in response to HIV-1 infection. The astrocyte production of IFN-β in response to various neurotropic viruses was documented (Pfefferkorn et al., 2016). Neither macrophages nor astrocytes produce type 1 IFN in significant amounts that are comparable to plasmocytoid dendritic cells seen in response to HIV-1 (Fitzgerald-Bocarsly et al., 2008; Wijewardana et al., 2009). We were able to confirm the possible source of IFN-β occurs through their interactions with infected macrophages. The transwell co-culture of HIV-1_ADA_-infected MDM and astrocytes revealed the temporally dependent dynamics of IFN-β production.

The cellular machinery that glia use to protect themselves from infection could also secondarily create an inflammatory cascade. This could then lead to restricted astrocyte infection and programmed cell death (Thompson et al., 2001). Through the use of unbiased image analysis we found a limited yet significant decrease in the number of human GFAP^+^ astrocytes in the brains of chimeric mice infected with HIV-1 and reduced expression of GFAP (Fig. 4 and 6). A similar scenario is known to occur in quiescent CD4 T cells, which undergo an abortive infection with accumulation of incomplete reverse viral transcripts (Monroe et al., 2014). In support of this theory, our data showed significant upregulation of genes serving as cytoplasmic sensors for nucleic acids, such as *DDX58, DDX60* and *IFI6* (Li et al., 2016). Therefore, the downstream signaling and effect of these RNA sensors in glial cells are certainly worthy of further investigation. We also cannot exclude compromised neurogenesis by HIV-1-induced interferon-related pathways, which preclude astrocyte’s differentiation (Carryl et al., 2015; Geffin and McCarthy, 2013).

The identified downregulated genes are mostly critical for oligodendrocyte differentiation and myelination. Among them, ZNF488 is a critical transcriptional factor that interacts and cooperates with Olig2 to promote oligodendrocyte differentiation (Wang et al., 2006). *PLP1, MAG, MOBP,* and *MBP* are encoding structure proteins of the myelin sheath. *CLDN11* encodes a crucial component of the tight junctions between the myelin sheath (Gow et al., 1999; Morita et al., 1999). Upregulation of LONGO1 expression also could inhibit myelination and oligodendrocyte differentiation (Jepson et al., 2012). Together, all of these transcriptional changes suggest early alterations in the structure and function of white matter during HIV infection. This may potentially lead to a compromise in the integrity of the glial system and consequent neural dysfunction.

In summary, we successfully established the dual reconstituted chimeric mouse model that contained both a human immune system and human glia. This model allowed us to access how glial-immunocyte crosstalk may be affected by HIV-1 infection. Using sequencing we found that the transcriptional profiles from the chimeric brains overlap, in large measure, with the disease profile found in human HIV-1 infected patients with cognitive impairments, encephalitic brains and acute SIV infection (Borjabad et al., 2011; Cserhati et al., 2015; Gelman et al., 2012; Sanfilippo et al., 2017; Sanna et al., 2017; Siangphoe and Archer, 2015; Winkler et al., 2012). These findings underscore the protective role astrocytes have in defending the brain from HIV-1 infection and damage of myelination processes. This model also provides a platform to further investigate the pathobiology of HIV nervous system infection and viral cell reservoirs needed to combat novel disease combating therapies and eradication measures.

## Materials and Methods

### Animals, cell transplantation, and HIV-1 infection

Newborn NOD/SCID/IL2Rγc^-/-^ (NSG, https://www.jax.org/strain/005557) mice were bred at the University of Nebraska Medical Center (UNMC). Animal procedures strictly followed the Institutional Animal Care and Use Committee approved protocols at UNMC and in adherence to the Animal Welfare Act and the Public Health Service Policy on Humane Care and Use of Laboratory Animals. P0-1 litters were irradiated with 1 Gy (RS 2000 X-ray Irradiator, Rad Source Technologies, Inc., Suwanee, GA, USA). Pups were intrahepatically injected with 5 × 10^5^ CD34^+^ HSPCs. These mice were simultaneously transplanted with 0.5 – 1×10^5^ NPC into the right lateral ventricle. Two samples of 97 gestation days (experiments 1 and 2) and two samples of 117 days gestation days (experiments 3 and 4) of liver and brain were used to create 4 cohorts of experimental animals. NPC were generated by 4 - 6 passages. Total 45 dual reconstituted animals were created, 23 females and 22 males. Blood samples collected from the facial vein were evaluated by flow cytometry starting 8 weeks post-engraftment to monitor expansion of human leukocytes. Mice with established human hemato-lymphoid reconstitution (∼5 months of age) were intraperitoneally infected (n = 22) with the macrophage-tropic ADA HIV-1 strain and euthanized 5 weeks after the infection was established (Table 1).

### NPC isolation and culture

Human NPC were isolated from human fetal brains provided by the University of Washington Medical Center Laboratory of Developmental Biology (R24HD000836-51). The UNMC Scientific Research Oversight Committee (SROC) and Institutional Review Boards at the University of Washington Medical Center and UNMC approved the protocol. Cells were isolated from human fetal brains as previously described (Nunes et al., 2003). Briefly, single cell suspensions were cultivated in NS-A medium (STEMCELL, Mountain View, Canada) containing N2 supplement (1:100; Fisher Scientific, Waltham, MA, USA), neural survival factor-1 (NSF1; 1:50; Lonza, Walkersville, MD, USA), LIF (10 ng/ml; Sigma-Aldrich, St. Louis, MO, USA), epidermal growth factor (EGF; 20 ng/ml; Fisher, MA, USA), and basic fibroblast growth factor (bFGF; 20 ng/ml; STEMCELL, Mountain View, Canada) for 14 days before passage. Four hours before injection, the neurospheres were dissociated into single cell suspension with 0.25% Trypsin (Gibco, Grand Island, NY, USA). The single cell suspensions were plated on polystyrene-coated coverslips and cultured for 48 hours in vitro. Immunofluorescent cytology was applied to investigate the differential potential of NPC.

### Human hematopoietic stem progenitor cells isolations

Matched to the brain sample, the fetal liver tissue was processed with 0.5 mg/ml DNase, collagenase, and hyaluronidase (all from Sigma-Aldrich, St. Louis, MO, USA) using gentleMACS(tm) Dissociator (Miltenyi Biotec, Auburn, CA, USA). Cell suspensions were passed through a 40-µm cell strainer and then centrifuged through leukocyte separation medium (MP Biomedicals, Santa Ana, CA, USA) at 100 g for 30 min, with acceleration set to 9, and deceleration set to 1. The intermediate layer of cells was collected for positive isolation of CD34^+^ cells using a magnetic isolation kit (Miltenyi Biotec, Auburn, CA, USA). CD34^+^ cells were stored in liquid nitrogen using a freezing medium with 50% fetal bovine serum (FBS), 40% Roswell Park Memorial Institute (RPMI)-1640 medium (Hyclone, Logan, UT, USA), and 10% dimethyl sulfoxide (DMSO; Sigma-Aldrich, St. Louis, MO, USA).

### Flow cytometry

Blood samples were collected from a facial vein in ethylenediaminetetraacetic acid (EDTA)-containing tubes (BD Microtainer, Franklin Lakes, NJ, USA) and centrifuged at 300 g for 5 min. Plasma was stored for HIV viral load measurements. Blood cells were reconstituted in a buffer of 2% FBS in phosphate buffered saline and incubated with antibodies against human cell markers, including CD45^+^ fluorescein isothiocyanate (FITC, BD/555482), CD3^+^ Alexa700 (BD/561805), CD19^+^ R-phycoerythrin (PE) cyanine 7 (Cy7, BD/555414), CD4^+^ allophycocyanin (APC, BD/ 561841), CD8^+^ Brilliant Violet^TM^ 450 (BV450, BD/562428), and CD14^+^ PE (BD/555398) for 30 min at 4°C. Samples were analyzed using a BD LSR2 flow cytometer using acquisition software FACS Diva v6 (BD Biosciences, USA), and data was analyzed using FLOWJO analysis software v10.2 (Tree Star, USA). Gates were assigned according to the appropriate control population.

### Measurements of viral load in plasma and brain tissues

Viral RNA copies in the murine plasma were determined by using a COBAS Amplicor System v1.5 kit (Roche Molecular Diagnostics, Pleasanton, CA, USA). Expression of HIV-1 group-specific antigen (gag) RNA in brain tissues were analyzed by reverse transcription (RT)-PCR with previously published primers and probes (Arainga et al., 2016). Briefly, cortex and striatum from three HIV-infected dual reconstituted mice and three control dual reconstituted mice were micro-dissected and transferred to 700 µl QIAzol solution (QIAGEN, Valencia, CA, USA) for RNA isolation using an RNeasy Mini kit (QIAGEN, Valencia, CA, USA). A Verso cDNA synthesis kit (Thermo Fisher Scientific, Grand Island, NY, USA) prepared cDNA according to the manufacturer’s instructions. For the first round of conventional PCR, we used the following primers: sense 5’ TCAGCCCAGAAGTAA-TACCCATGT 3’ and antisense 5’ TGCTATGTCAGTTCCCCTTGGTTCTCT 3’. The PCR settings were as follows: 94°C for 3□min, 15 cycles of 94°C for 30□sec, 55°C for 30□sec, and 72°C for 1□min. The product of the first PCR was subsequently used as a template in the second semi-nested real-time PCR amplification performed on the ABI StepOne Plus real-time PCR machine (Applied Biosystems, MA, USA) using TaqMan detection chemistry. The primers and probe used for the second PCR were: sense 5’ TCAGCCCAGAAGTAATACCCATGT 3’, antisense 5’ CACTGTGTTTAGCATGGTGTTT 3’, and TaqMan probe FAM-ATTATCAGAAGGAGCCACCCCACAAGA-TAMRA. The real-time PCR settings were as follows: 50°C for 2□min, 95°C for 10□min, 45 cycles of 95°C for 15 sec, and 60°C for 1□min. ACH2 cells (8□×□10^5^) containing one integrated copy of HIV-1 per cell were used in triplicate as standards, with cell and HIV copy numbers ranging in serial 10-fold dilutions from 10^5^ to 10^2^ deoxyribonucleic acid (DNA) copies/reaction.

### Immunohistochemistry

Brain tissue was cut sagittally and the left and right hemispheres from experiments 1 and 2, and right hemispheres from experiments 3 and 4, were fixed with 4% paraformaldehyde for 24 hours at 4°C, and then embedded in paraffin. Paraffin-embedded 5-µ thick tissue sections were processed with Declere Solution (Sigma-Aldrich, St. Louis, MO, USA) according to the manufacturer’s instructions. Tissue was blocked with 10% normal goat serum in Tris-buffered saline and 0.05% Tween 20 (TBST) for 45 min and then incubated with mouse (ms) anti-human GFAP (1:1000; Y40420/STEM123, TaKaRa Bio, USA), ms anti-human nuclear antigen (1:200; Millipore/MAB4383, Billerica, MA, USA), ms anti-human cytoplasmic marker (1:1000; Takara/STEM121, Mountain View, Canada), ms anti-Nef (1:200, Santa Cruz/sc-65904), ms anti-HLA-DR (1:500; Dako/M0746, Carpinteria, CA, USA), ms anti-CD163 (1:50; Thermo/MA5-11458, Denver, CO, USA), rabbit (rb) anti-GFAP (1:1000; Dako/Z0034, Carpinteria, CA, USA), and rb anti-human CD4 (1:500; Abcam/ab133616). Secondary antibodies were goat anti-ms and goat anti-rb immunoglobulin G (IgG) horseradish peroxidase (HRP; Dako, Carpinteria, CA, USA). Bright field images were captured and photographed using 20× and 100× objectives on a Nuance Multispectral Tissue Imaging system (CRi, Wobum, MA). Immunostained sections were scanned using a high-resolution scanner (Ventana Medical Systems, Inc., Oro Valley, AZ, USA). DEFINIENS Tissue Studio**®** software (Definiens AG, Munich, Germany) was used to analyze the brain sections stained for human glial and immune cells. Spleen and brain tissue sections were analyzed for HIV-1 p24 protein staining with anti-HIV-1 p24 (1:10; Dako/M0857, Carpinteria, CA, USA). None of human-specific antibodies had cross-reactivity and stained mouse brain sections not reconstituted with human cells.

### Immunofluorescence

For immunofluorescent staining of paraffin-embedded tissue, sections were processed and blocked as described above. Ms anti-human GFAP, ms anti-human nuclei, rb anti-GFAP, and rb anti-Oligo2 (1:500; PhosphoSolution/1538, Aurora, CO, USA) were applied. A Leica DM6 system (Leica Microsystems, Wetzlar, Germany) were used for immunofluorescent imaging. The intensity of illumination and the position of the sub-region on the microscope were consistent across all images. For immunofluorescent characterization of cultivated NPC, ms anti-human GFAP (1:200; Abcam, Cambridge, MA, USA), ms anti-MAP2 (1:200; Santa Cruz, Santa Cruz, CA, USA), ms anti-glutamic acid decarboxylase 67 (GAD67; 1:500; Millipore, Billerica, MA, USA), rb anti-nestin (1:200; Millpore, Billerica, MA, USA), and rb anti-Tuj-1 (1:500; Biolegend, San Diego, CA, USA) were applied. Secondary antibodies were goat anti-ms IgG Alexa 488 (1:00; Invitrogen, Grand Island, NY, USA), goat anti-ms IgG Alexa 594 (1:00; Invitrogen, Grand Island, NY, USA), goat anti-rb IgG Alexa 488 (1:100; Invitrogen, Grand Island, NY, USA), and goat anti-ms IgG Alexa 594 (1:00; Invitrogen, Grand Island, NY, USA). A Leica DM6 system (Leica Microsystems, Wetzlar, Germany) and a Zeiss LSM510 confocal system (Carl Zeiss Microscopy GmbH, Jena, Germany) were also used for immunofluorescent imaging.

### Next generation sequencing

Left hemispheres from experiments 3 and 4 of the reconstituted mouse brains (0.5 × 10^5^ cells/mouse) were dissected by regions and flash-frozen for RNA extraction. The brain of three age-matched NSG mice were also collected for non-reconstituted control. For sequencing hippocampus (Hip) and corpus callosum (CC) RNA from brain tissue from three uninfected dual-reconstituted animals, 3 HIV-1 infected mice and 3 NSG non-reconstituted controls were preserved (Invitrogen, Grand Island, NY, USA). Tissue was transferred to 700 µl of QIAzol solution (QIAGEN, Valencia, CA, USA) for RNA isolation using an RNeasy Mini kit (QIAGEN, Valencia, CA, USA). After nucleic acid integrity was analyzed, samples were deep sequenced using 100 bp/read, up to 40 million reads/sample using an Illumina HiSeq 2500 Sequence Analyzer (Illumina, Inc., San Diego, CA, USA). Reads were trimmed using the fqtrim software to remove ambiguous bases from the reads. Read alignment and differential gene expression was performed using the programs from the Tuxedo RNA-seq tool pipeline. The trimmed reads were then aligned to the mouse genome (version mm10/GRCm38) and the hg19 human genome (when necessary) using the tophat aligner, version 2.1.1. The ratio of aligned reads per total reads was calculated to assure quality alignments. Unaligned reads coming from the humanized samples were remapped to the hg19 database. FPKM values (Fragments Per Kilobase of transcript per Million mapped reads) were calculated for each gene in each sample using the cufflinks software, version 2.2.1. This was done to calculate each gene’s expression level in each sample. An FDR adjusted p-value of 5% was used in all cuffdiff sample comparisons. For both regions (Hip and CC) the control samples (NSG unmanipulated) to the humanized mouse samples (blood and brain reconstituted), and the humanized mouse samples to the humanized mouse samples with HIV were compared. The reads were mapped to the mm10 mouse genome and the mouse transcriptome. For both tissues, the unmapped reads, which did not map to the mouse genome, were collected and remapped to the human genome. This was done in order to see what human genes were expressed in these tissues. Cuffdiff was specifically used because it normalizes the expression values and protects against overdispersion so that the values fit into a normalized bell curve (Trapnell et al., 2012).

### Macrophage and human fetal astrocytes isolation and cultivation

Human monocytes were isolated by leukopheresis from HIV-1/2 and hepatitis seronegative donors. Human monocytes (2 × 10^5^) were seeded in the inserts of 24-well transwell plates with a pore size of 0.4 µm. (Corning Costar, Corning, NY, USA). Dulbecco’s Modified Eagle’s Medium (DMEM; Invitrogen, Grand Island, NY, USA) with 10% heat-inactivated pooled human serum, 2 mM L-glutamine, 50 µg/ml gentamicin, 10 µg/ml ciprofloxacin, and macrophage colony-stimulating factor (MCSF) enriched conditioned medium maintained at 37°C in a 5% CO_2_ incubator was used (Gendelman et al., 1988). On day 7, differentiated monocyte-derived macrophages (MDM) were infected with HIV-1_ADA_ (multiplicity of infection (MOI) = 0.01) for 12 hours. Infected macrophages were maintained in the DMEM medium containing 10% heat-inactivated pooled human serum, 2 mM L-glutamine, 50 µg/ml gentamicin, and 10 µg/ml ciprofloxacin for another 6 days before the inserts were transferred to wells with astrocytes at the bottom. Human fetal astrocytes (HFA) were isolated and cultured from fetal donors (Nath et al., 1995). HFA (5×10^5^) were seeded at the bottom of 24-well transwell plates and maintained in DMEM/F12 (Invitrogen, Grand Island, NY, USA) containing 10% FBS and antibiotics. Supernatants from HIV-1-infected and control MDM cultures were collected from 2 to 6 days before MDM and astrocyte co-cultivation. Supernatants from the transwell system were collected at 24 hours, 3, 5, and 7 days.

### IFN-β enzyme-linked immunosorbent assay

IFN-β was measured in the supernatants of astrocytes, a co-culture of astrocytes and control macrophages, a co-culture of astrocytes and HIV-infected macrophages, HIV-infected macrophages, and control macrophages using the Human IFN-β bioluminescent ELISA kit (http://www.invivogen.com/lumikine-hifnb) according to the manufacturer’s instructions.

### RT-PCR for human and mouse transcripts

First-strand cDNA was prepared from CC and Hip using the same method described above. For both brain regions, we compared the control humanized mouse samples to the humanized mouse samples with HIV. The catalog numbers of all TagMan® real-time PCR assays from Life Technologies are as listed below: hu-STAT1 (Hs01013996_m1), hu-STAT2 (Hs01013123_m1), hu-IRF9 (Hs00196051_m1), hu-MX1 (Hs00895609_m1), hu-ISG15 (Hs01921425_s1), hu-CMPK2 (Hs01013364_m1), hu-IFI6 (Hs00242571_m1), hu-MBP (Hs00921945_m1), hu-ZNF488 (Hs00289045_s1), hu-GAPDH (Hs03929097_g1), ms-MX1 (Mm00487796_m1), ms-MBP (Mm01266402_m1), ms-GAPDH (Mm999999_g1). The real-time PCR settings were as follows: 50°C for 2□min, 95°C for 10□min, 40 cycles of 95°C for 15 sec, and 60°C for 1□min. All real-time assays were carried out with an Applied Biosystems® StepOnePlus Real-Time PCR Systems. The fold change of each target gene mRNA relative to GAPDH between the HIV^+^ group and the control group was determined. This was done by using threshold cycle (Ct) and the 2^-ΔΔ*C*^T method, where ΔCt = Ct_target_ – Ct_GAPDH_ and Δ(ΔCt) =ΔCt_HIV_ – ΔCt_control_.

### Statistical analysis

Values from all quantified astrocytes, oligodendrocytes, macrophages and lymphocytes and cell groups were averaged and presented as mean ± SEM; multiple t-tests using the Holm-Sidak method for multiple comparisons correction was applied to examine significant differences between groups. Two-way analysis of variance (ANOVA) using Sidak’s multiple comparisons correction was used to examine significant differences between groups. For RNA sequencing data, the tophat/cuffdiff pipeline was run on the reads for each sample in order to identify differentially expressed genes (Trapnell et al., 2012).

## Acknowledgements

The authors thank Prasanta Dash, Raghubendra Sing Dagur and Mariluz Anamelva Arainga Ramirez for technical support. We thank Li Wu, Yan Cheng, and Amanda Branch-Woods for assistance in preparing cells and managing animals. Assistance form Cathy Gebhart, Technical Director of The Molecular Diagnostics Laboratory, for viral load analysis is noted and appreciated. We acknowledge the UNMC Flow Cytometry Research Facility (Philip Hexley, Director) for technical support.

### Competing interests

‘No competing interests declared’

### Funding

We acknowledge support through the Office of the Vice Chancellor for Research and by state funds from the Nebraska Research Initiative (NRI) and The Fred and Pamela Buffett Cancer Center's National Cancer Institute Cancer Support Grant. Major instrumentation was provided by the Nebraska Banker's Fund and by the National Institute of Health (NIH)-National Center for Research Resources (NCRR) Shared Instrument Program. The University of Nebraska DNA Sequencing Core (James Eudy, Director) receives partial support from the National Institute for General Medical Sciences (NIGMS) Institutional Development Award Networks of Biomedical Research Excellence (INBRE; P20GM103427-14) and Centers of Biomedical Research Excellence (COBRE; 1P30GM110768-01) grants, as well as the Fred & Pamela Buffett Cancer Center (P30CA036727). We thank the Bioinformatics and Systems Biology Core Facility (Chittibabu (Babu) Guda, Director, and Matyas F. Cserhati) for providing data analysis services. This facility receives support from the Nebraska Research Initiative (NRI) and the NIH (2P20GM103427 and 5P30CA036727). This work was supported, in part, by the Dr. Carol Swarts Emerging Neuroscience Research Laboratory and the Frances and Louie Blumkin Foundation. Direct support from ViiV Healthcare, and NIH grants (P01 DA028555, R01 NS36126, P01 NS31492, 2R01 NS034239, P01 MH64570, P30 MH062261, R01 AG043540 all to HEG, R24 OD 018546 to LYP & SG, and 1R21 DA041018 to LYP & SG).

